# Electroencephalogram Monitoring of Depth of Anesthesia during Office-Based Anesthesia

**DOI:** 10.1101/2020.10.27.356592

**Authors:** Sunil B. Nagaraj, Pegah Kahali, Patrick L. Purdon, Fred E. Shapiro, M. Brandon Westover

## Abstract

**Objective:** Electroencephalogram (EEG) monitors are often used to monitor depth of general anesthesia. EEG monitoring is less well developed for lighter levels of anesthesia. Here we present an automated method to monitor the depth of anesthesia for office based procedures using EEG spectral features.

**Methods:** We analyze EEG recordings from 30 patients undergoing sedation using a multimodal anesthesia strategy. Level of sedation during the procedure is coded using the Richmond Agitation and Sedation Scale (RASS). The power spectrum from the frontal EEG is used to infer the level of sedation, by training a logistic regression model with elastic net regularization. Area under the receiver operator characteristic curve (AUC) is used to evaluate how well the automated system distinguishes awake from sedated EEG epochs.

**Results:** EEG power spectral characteristics vary systematically and consistently across patients with the levels of light anesthesia and relatively healthy patients encountered during office-based anesthesia procedures. The logistic regression model using spectral EEG features distinguishes awake and sedated states with an AUC of 0.85 (± 0.14).

**Conclusions:** Our results demonstrate that frontal EEG spectral features can reliably monitor sedation levels during office based anesthesia.

## INTRODUCTION

As surgical and anesthetic techniques advance, procedures are increasingly performed in ambulatory settings (Urman et al. 2012; Shapiro et al. 2014). Approximately one-third of ambulatory or “office based” anesthesia (OBA) services in the United States are provided as Monitored Anesthesia Care (MAC) (Bayman et al. 2011). During MAC, the anesthesia care team continually assesses the level of sedation to avoid an unintended progression to a state of general anesthesia (Shapiro et al. 2014). Nevertheless, oversedation leading to respiratory depression is still an important mechanism of patient injuries during MAC (Purdon et al. 2015). Incurrent OBA practice, sedation levels are either indirectly monitored through clinical observation, or based on evaluations of a patient’s responsiveness to verbal or tactile stimuli (Sheahan and Mathews 2014). However, the frequency and consistency of such evaluations is subjective and can be flawed (Green and Mason 2010). Moreover, effective drug doses to achieve a certain level of sedation in OBA may vary markedly between individuals (Shapiro et al. 2014; Kim et al. 2017). Therefore, implementing explicit and objective methods to assess sedation levels may improve safety in OBA practice.

Tracking brain state using EEG is a principled physiology-based method to achieve reliable continuous assessment of OBA sedation levels. Previous studies have found that using EEG-based monitoing during procedural sedation is accompanied by reduction in propofol administration (Conway and Sutherland 2016; Park et al. 2016), a commonly used intravenous hypnotic agent for ambulatory anesthesia (Seamans 2008; Akeju et al. 2016). However, the EEG response can be variable due to the use a multimodal apporach (i.e. using several drugs in combination) that targets specific endpoints including hypnosis, amnesia, analgesia and akinesia (Brown et al. 2010). Multimodal anesthesia involves the use of smaller doses of several drugs with the goal of maximizing the benefits and minimizing the side effects of each drug.

The objective in this study is to evaluate the feasibility of automated EEG-based sedation level tracking using the EEG power spectrum during multimodal anesthesia in the office based setting.

## METHODS AND MATERIALS

### Patient selection

The study was performed with under a protocol approved by the ethics committees of Beth Israel Deaconess Hospital and Massachusetts General Hospital, Boston, USA. Waiver of consent was granted to analyze EEG recorded during general anesthesia or sedation. The EEG recordings were strictly observational and were not used for clinical management. In total, we obtained EEG recordings from 46 patients undergoing multimodal sedation for either esophagogastroduodenoscopy or colonoscopy or both, for diagnostic and therapeutic purposes, during the period September 2016-March 2017. We excluded 16 patients from the analysis due to technical problems during EEG recording. Therefore, we analyzed the data from 30 patients (16 males, 14 females, mean age: 60.8±12.9, mean weight = 78.8±18.4 kg, mean height =168.6±13.1 cm) in this study.

### EEG recording

We used SedLine 4-channel forehead EEG monitors (Massimo Corporation, Irvine, 40 CA) to record EEG signals from patients undergoing OBA sedation. We placed recording electrodes on each patient’s forehead approximately at positions Fp1, Fp2, F7, and F8, 1 cm above the ground electrode FpZ. We ensured that the impedances in each channel were < 5kΩ before EEG recording. We did not adjust the electrodes for impedance after starting the surgical procedure. We used the following settings to record EEG: sampling frequency = 250 Hz, preamplifier bandwidth = 0.5-90 Hz and resolution = 16-bit, 29 nV.

### Sedation assessment

For scoring patients’ level of sedation we used the RASS behavioral scale (Sessler et al. 2002). RASS scores range between −5 (unresponsive to external stimuli) to 0 (awake/calm). Higher numbers (+1 through +4), not used in our study, denote levels of agitation. Research staff (*MBW, SBN, PK*) assigned RASS scores approximately once every 30s throughout the procedure. Clinical staff received training to reduce inter-rater variability in RASS assessments. For training RASS prediction models, we group EEG epochs with RASS scores = 0 and −1 (“awake”), and epochs with RASS scores = −4, −5 (“sedated”); epochs with intermediate scores (−3, −2) are not used for model training. This ensures that the data used to train the classification model represent clearly distinguishable clinical states.

### Anesthesia Protocol

The anesthesia care team reviewed patients’ medical histories, including pre-existing medical conditions, drug allergies, medications, social history and medical records to select appropriate drugs. MAC anesthesia was used for all patients. Induction and maintenance of sedation was performed using a continuous low-dose infusion of propofol (25-75 μg/kg/minute), with intermittent boluses of propofol (20-100 mg), ketamine (20-40 mg), dexmedetomidine (1 μg/kg over 10 minutes) and lidocaine (5-10 mg) (Shapiro 2007). In several cases the anesthesia provider modified the infusion rate based on the patients’ assessed level of sedation. A few patients were pre-medicated with a single dose of haloperidol (0.25 mg) and midazolam (1-2 mg) if they displayed increased anxiety in the preoperative holding area. All patients were monitored according to the Standards for Basic Anesthetic Monitoring established by the American Society for Anesthesiologists (ASA), which requires the continuous evaluation for oxygenation, ventilation, circulation and temperature during all anesthetics. Changes in drug infusion rates and timing and doses of drug boluses throughout the procedure were recorded using logging software. The anesthesia provider delivered care independent of EEG monitoring.

## AUTOMATIC SEDATION LEVEL MONITORING SYSTEM

The architecture of the proposed EEG-based automatic sedation level monitoring system is shown in Figure 1. Details of each stage are described below.

**Figure 1:**
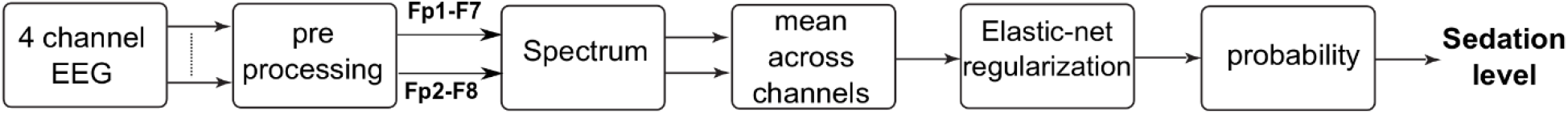
Architecture of the proposed automatic sedation level detection system.

### Preprocessing and artifact rejection

We use a bipolar montage, with channels defined by the differences Fp1-F7 and Fp2-F8. The EEG signal is passed through a bandpass filter set at 0.5-25 Hz. For model training, frequency range is limited to 0.5-25 Hz to reduce the influence of muscle artifact during the awake state. Using a 5 second moving window we detect artifacts in the EEG signal based on the following conditions: (1) amplifier saturation/movement artifacts – abnormally high signal amplitude (> 500 *μV*); (2) 60 Hz activity above 500 *μV*(measured spectrographically); (3) loose electrode artifacts – mean amplitude of the sum of signals less than half the mean amplitude of the first channel. We segment the EEG into overlapping 4s epochs with 0.1s shift for further analysis.

### Feature extraction

We perform spectral estimation using multitaper spectral analysis via the chronux toolbox (Bokil et al. 2010) with the following parameters: length of the window T = 4s with 0.1s shift, time-bandwidth product TW = 3, number of tapers K = 5, and spectral resolution 2W of 1.5 Hz. Figure 2 shows an example of the spectrogram along with RASS assessments and drug infusion/bolus rates from a single subject.

**Figure 2:**
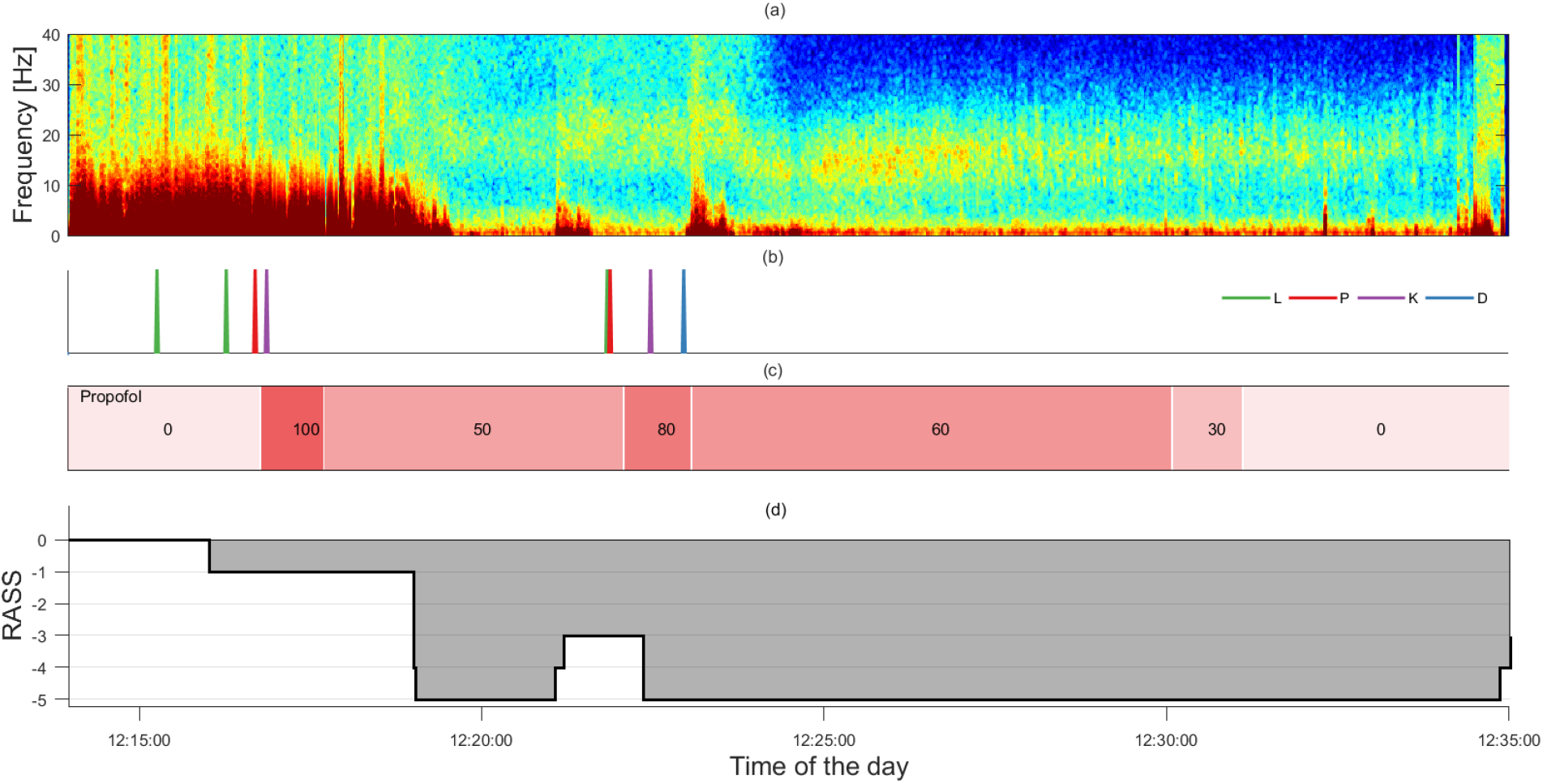
**Sample figure from a patient showing (a) Frontal EEG multitaper spectrogram, (b) bolus dosages: L = Lidocaine, P = Propofol, K = Ketamine, and D = Dexmedatomidine, (c) continuous propofol infusion rate, and (d) RASS assessments.**

We use both the un-normalized (raw) power spectrum and normalized power spectrum (normalized by total power) averaged across the two bipolar montages to train the logistic regression model. Figure 3 shows the EEG signal and corresponding spectrogram at different RASS levels in one subject.

**Figure 3:**
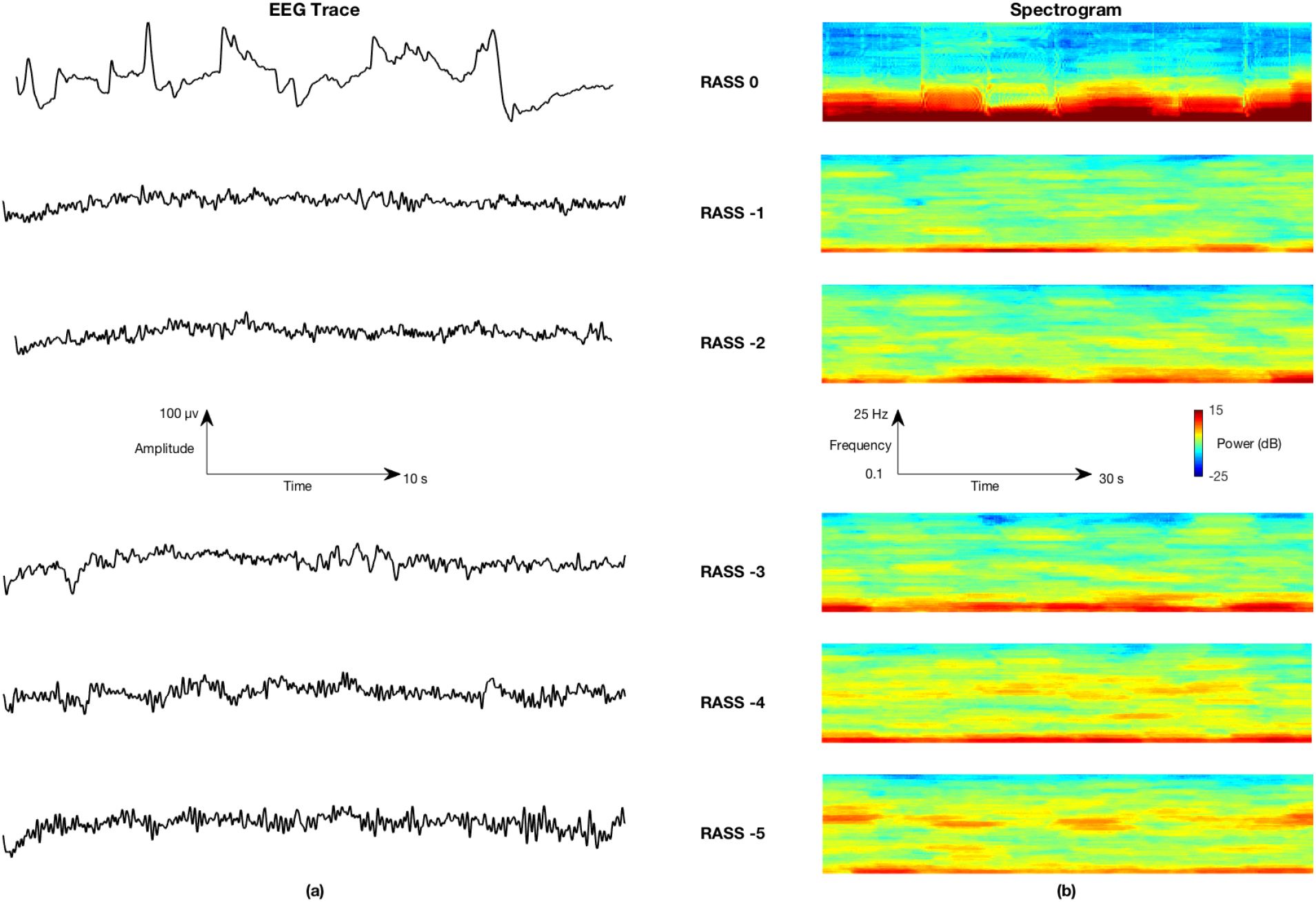
(a) Example of EEG signal at different RASS states and (b) their corresponding multitaper spectrogram.

### Classification and post-processing

We use logistic regression with with elastic-net regularization (Tibshirani 1996) to train models to detect sedation levels. The benefit of using elastic-net regularization is that it has low computational complexity and is suitable for for high-dimensional data with multiple correlated features (Efron et al. 2004). In addition, by setting many of the coefficients to zero, elastic-net automatically performs feature selection.

### Model Performance Evaluation

#### Metrics

We use the area under the receiver operator characteristic curve (*AUC*) as a metric to assess the performance of the model for binary classification. In addition, we use the Spearman rank correlation coefficient (*ρ*) to measure the correlation between the probability output of the model with RASS scores over time. The rationale is that *ρ* provides information complementary to AUC: it measures the extent to which increases or decreases in sedation depth are tracked by changes in the model output, inclluding for intermeiate RASS values. Thus is a measure of how well the model performs in providing a continuous output, as as opposed its ability to provide binary predictions.

#### Cross validation

We use leave-one-patient-out (LOPO) cross-validation to assess performance of the prediction system. In this method, data from *N*-1 patients are used for training the system and ata from the one remaining patient is used for testing. This is repeated until each patient’s data is used for testing, resulting in a total of *N* iterations. During training, we standardize the training set features to have unit mean and standard deviation. We also normalize features in the testing with respect to the mean and standard deviation of the training set so that all data is processed in the same way. To optimize the set of features to be included in the final model (“feature selection”), and model parameters (*λ*, *α*), required by the elastic-net algorithm, we use 10-fold cross validation on the training set to obtain these parameters. We use the parameters (*λ*, *α*) that provide maximum mean AUC over 10-folds of training data to train the final model on all the training data. After obtaining the final trained model with optimal parameters in the inner cross validation loop, we obtain the probability of sedation on the left-out testing subject in the the outer cross validation loop of LOPOCV (30 iterations in total). In this way, we only use training data to optimize model parameters. The model is kept independent of the testing data, mimicing the real clinical scenario.

#### Training and testing strategy

We evaluate two different training and testing combinations: 1) TBTB – train and test on binary data in which the model is trained and tested to discriminate between awake and sedated epochs, and 2) TBTR – train on binary data and test on RASS scores (ordinal RASS −5,-4,…,0 scores rather than binarized scores). In this evaluation, we use the binary classifier which is trained on awake and sedated epochs to assign a probability score to *all* EEG epochs with RASS scores. We calculate the Spearman rank correlation coefficient (*ρ*) between the probability score and the RASS scores. We perform all coding and analysis using the MATLAB 2018a scripting language (Natick, USA).

## RESULTS

Unless stated otherwise, all results are reported as mean (± SD).

### Automatic sedation level classification

#### (i) TBTB – Binary classification

Figure 4 shows the coefficient weights assigned by the elastic-net model as a function of frequency. The oscillatory activity around the slow-delta (0.1-4 Hz) band shows negative correlation with the awake state (RASS = 0 and −1), as opposed to the oscillatory activity around high alpha and low beta (9-15 Hz) bands, which correlates with the sedated state (RASS = −4 and −5). The multivariate logistic regression model using spectrogram (normalized + un-normalized spectrogram) resulted in an AUC = 0.85 (0.14).

**Figure 4:**
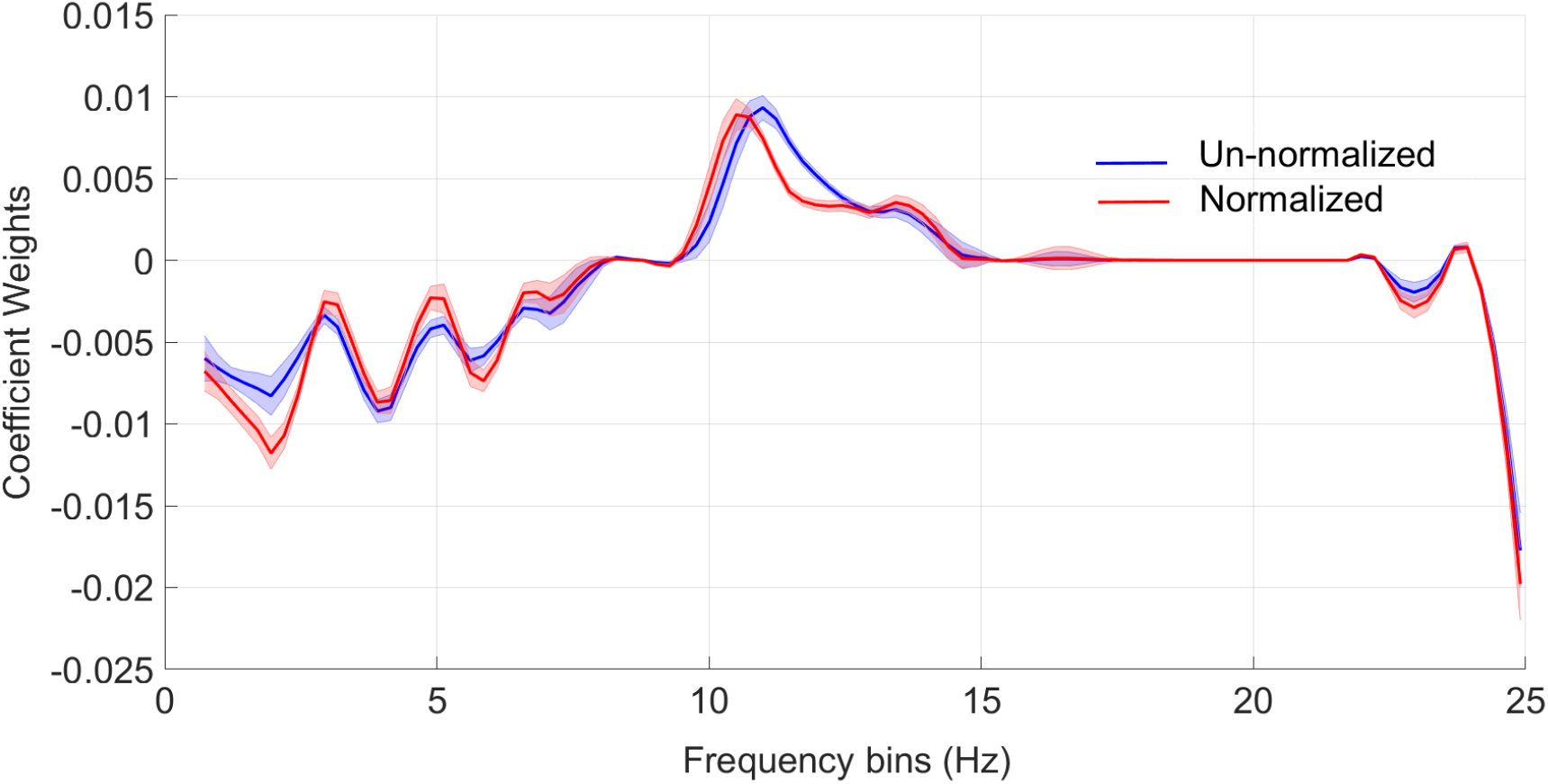
Plot of elastic-net regression coefficients of all frequency bins. The regression coefficients are estimated from the elastic net analysis of the un-normalized (blue) and the normalized spectrogram (red) across all subjects.

#### (i) TBTR – Correlation with continuous RASS scores

Although during training we optimized the system to make a binary distinction between sedated (RASS = −5 and −4) and awake (RASS = 0 and −1) states, the model provides a continuous output, i.e. a probability of being awake. To investigate the potential utility of this probability as an index of depth of sedation during OBA, we next evaluate the correlation between the model output over time and all RASS states between − 5 to 0. This results in a mean *ρ =* 0.38 (0.17) across 30 patients, substantially better than chance level correlation (*ρ =* 0.18 (0.11)). This suggests that the automated system, despite being trained to make binary discriminations, provides significant information about sedation levels along the continuum between sedation and wakefulness. An example illustrating this process is shown in figure 5.

**Figure 5:**
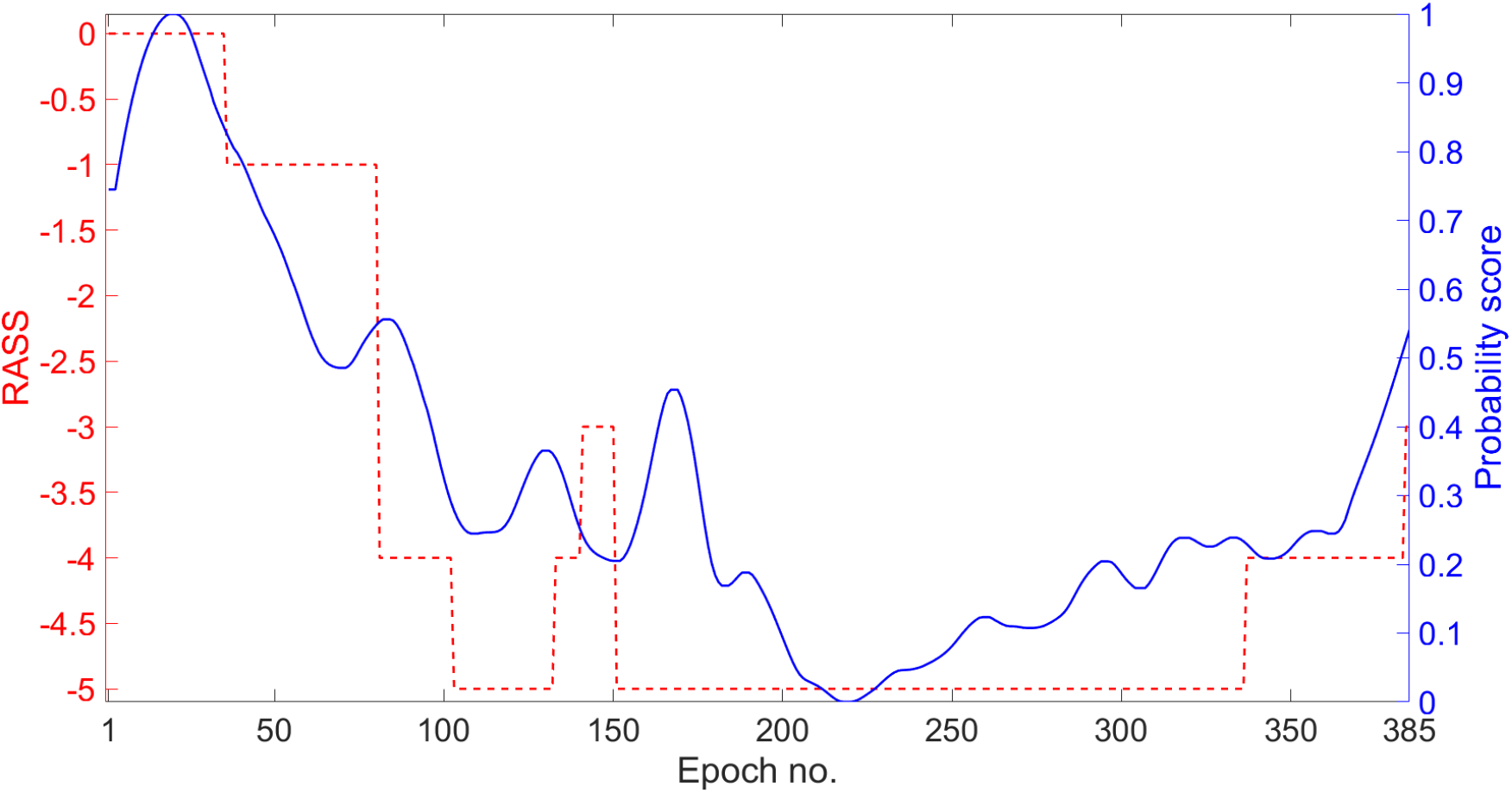
**Example of the correlation between the probability score of the classifier output with RASS assessments (*ρ* = 0.68 in this example). In this method, a trained binary linear model (trained only on awake and sedated epochs) is used to obtain continuous probability score on the testing patient.**

## DISCUSSION

In this study, we investigated the potential of a neurophysiology based system to characterize sedation levels, using spectral features of the frontal EEG, in a multimodal office-based anesthesia setting. Our results show that slow-delta (0.1-4 Hz) oscillations are associated with lighter sedation levels (RASS = 0 and −1) induced by propofol when combined with adjunctive sedatives and analgesics. We also observed that high alpha-low beta (9-15 Hz) oscillations are associated with deeper levels of sedation (RASS =−4 and −5). Our results show that spectral features of the EEG show systematic trends with increasing levels of sedation that computer algorithms can leverage to infer the level of sedation. Using statistical learning we identified weighted combinations of spectral EEG features that provide good discrimination between awake and sedated EEG states (AUC = 0.85 (0.14)). Our system provides a continuous probability estimate of the patient’s sedation level.

Monitoring depth of sedation during standard anesthesia care is typically based on behavioral assessments of the patient's arousability in response to verbal or tactile stimuli and indirectly through monitoring cardiovascular and respiratory function, whereas intermittency in behavioral assessments and subjectivity in identifying and quantifying a patient’s response (Green and Mason 2010) can result in inadequate sedation or oversedation. EEG-based anesthesia monitors are often used when general anesthesia is required. However, explicit and objective brain monitoring may also benefit cases targeting lighter sedation levels, to prevent unintended deeper levels of anesthesia.

The multimodal sedation strategy is gaining popularity, particularly in ambulatory settings, due to better safety profiles. The utility of EEG-based monitors has been less investigated when anesthetics are administered concurrently with other hypnotics, sedatives and analgesics. Concurrent administration of centrally-acting hypnotics, sedatives and analgesics induce anesthetic states that might have different neurophysiologic properties from those when single anesthetics are used. Therefore, it is important to investigate how well EEG based monitoring of sedation levels performs in the setting of multimodal sedation.

## LIMITATIONS

There are several limitations in this study. First, we only explored features derived from the EEG spectrogram. This has the advantage of straightforward interpretation, but it is possible that less interpretable features improve model performance. Future work will explore whether additional features (entropy, complexity) can improve the performance of the proposed system. A second limitation is the limited number of patients. Future work will need to validate the robustness of the system on a dataset from a large cohort of patients undergoing multi-drug sedation. Third, we used a simple logistic regression approach to train the sedation level prediction model. More advanced machine learning models may improve the performance of the proposed system. Finally, we only trained the model to perform binary classification in this study (awake vs sedated). Future work will explore discriminating multiple sedation levels on a continuous scale using a prospective dataset.

## CONCLUSION

This study evalutes the performance of an automated system using the EEG spectrogram for predicting sedation levels in an office based anesthesia setting, using a multimodal sedation strategy. Our findings suggest that features from the EEG spectrogram are able to discriminate awake vs sedated brain states.

## DATA AVAILABILITY

The data used in this study is available from github repository: https://github.com/mghcdac/EEG_mulitmodal_sedation

